# Relevance of genomic evaluation for egg quality traits in layers

**DOI:** 10.1101/704742

**Authors:** David Picard Druet, Amandine Varenne, Florian Herry, Frédéric Hérault, Sophie Allais, Thierry Burlot, Pascale Le Roy

## Abstract

**Background:** Genomic evaluation, based on thousands of genetic markers, has become the standard evaluation methodology in dairy cattle breeding programs over the past few years. Despite the many differences between dairy cattle breeding and poultry breeding, genomic selection seems very promising for the avian sector, and studies are currently being conducted to optimize avian selection schemes. In this optimization perspective, one of the key parameters is to properly predict the accuracy of genomic evaluation in pure line layers.

**Methods:** Both genetic evaluation and genomic evaluation were performed on three candidate populations (male and female), using different sizes of phenotypic records on five egg quality traits and at two different ages. The methodologies used were BLUP & ssGBLUP, and variance-covariance matrices were estimated through REML. To estimate evaluation accuracy, the LR method was implemented. Four statistics were used to assess the relative accuracy of the estimated breeding values of candidates, their bias and dispersion, as well as the differences between genetic evaluation and genomic evaluation.

**Results:** It was observed that genomic evaluation, whether performed on males or females, always proved more accurate than genetic evaluation. The gain was higher when phenotypic information was narrowed and an augmentation of the size of the reference population led to an increase in accuracy prediction, for what regards genomic evaluation. By taking into account the increase of selection intensity and the decrease of the generation interval induced by genomic selection, the expected annual genetic gain would be higher with ancestry-based genomic evaluation of male candidates than with genetic evaluation based on collaterals. This advantage of genomic selection over genetic selection requires to be studied in more details for female candidates.

**Conclusions:** In conclusion, in the population studied, genomic evaluation for egg quality traits of breeding birds at birth seems a promising strategy, at least for what regards males selection.

## Introduction

Genomic evaluation, based on thousands of genetic markers [1], has become the standard evaluation methodology in dairy cattle breeding programs over the past few years. It has allowed for the improvement of the accuracy of estimated breeding values (EBV) of young animals and for the reduction of the generation interval, as well as for the reduction of the phenotyping costs [2]. More recently, avian breeders have started to implement genomic selection in their selection schemes. Indeed, despite the many differences between dairy cattle breeding and poultry breeding, genomic selection is deemed very promising for the avian sector, especially for layers selection [3, 4, 5]. However, to optimize avian selection schemes, one of the key parameters is to properly predict the accuracy of genomic evaluation.

One of the most important factors directly affecting evaluation accuracy is the makeup of the reference population. From the very beginning, genomic evaluation implied that the size of the reference population should not be too small [1, 2, 6]. However, it has also been shown that increasing the size of the reference population does not directly improve evaluation accuracy [7, 8]. Indeed, more than the size of the reference population, it is its propinquity with the candidate population that is critical. The evaluation is all the more accurate as candidates haplotypes are well represented in the reference population [7, 9, 10, 11, 12]. Aside from the makeup of the reference population, the number of training generations to use is another important question. Indeed, it has been shown that evaluation accuracy is impacted by the number of training generations used [13, 14], depending on the heritability of the traits.

The present study assesses the relevance of genomic evaluation in comparison with genetic evaluation, in order to predict the breeding values of selection candidates for egg quality traits in a pure line of layers. The main objective was to evaluate the expected genetic gain on those traits, in order to move from genetic to genomic evaluation.

## Material and Methods

### Animals

For the purpose of this study, we have used the data of a pure line of Rhode Island layers selected by the breeding company Novogen (Plédran, France). The hens were hatched in 12 batches, born between 2008 and 2015, which corresponds to four generations (G0 to G3, cf figure 1), with three successive hatches per generation spaced 6 months apart. The genealogy of all the birds was recorded in the pedigree file, which concerned 2,273 breeders: 514 sires and 1,759 dams.

**Figure. 1.**
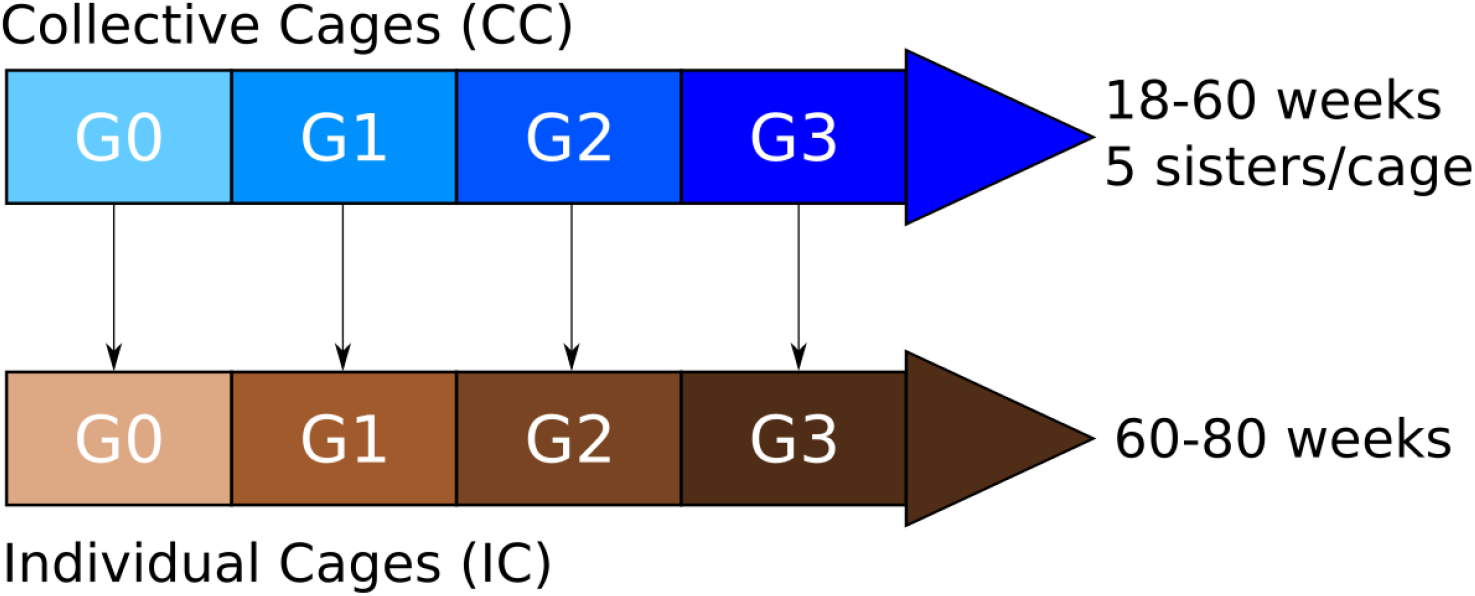
Population structure.

**Figure. 2.**
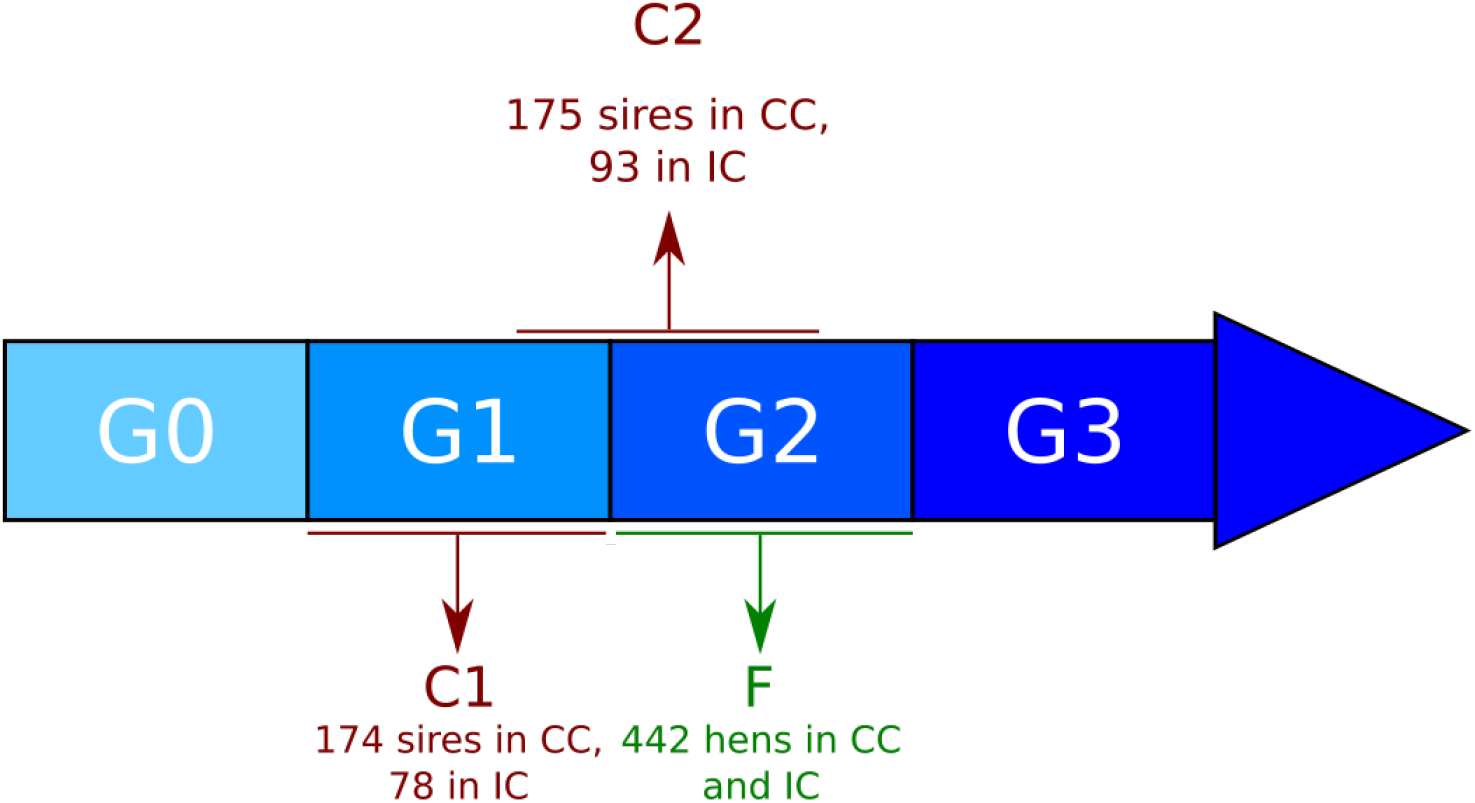
Candidates populations.

In each hatch, chicks were bred in a brooding area until the age of 18 weeks, and then transferred to collective cages of five full sisters for females (2,997 collective cages) and to individual cages for males (200 males out of 2,000 chicks were kept for the selection). Egg quality was recorded twice during this period, at 30 and 50 weeks of age. This step allowed for the makeup of our first phenotypic data set, referred to as CC for collective cages, with a total of 14,985 hens and 27,915 eggs measured.

Then, at 60 weeks of age, a genetic evaluation was performed as a first selection, and 150 males and 600 hens were transferred to individual cages, until the end of their career, at the age of 80 weeks. Egg quality was measured on a weekly basis. A total of 7,982 hens, with 74,976 performances, were concerned. This step allowed for the makeup of our second phenotypic data set, referred to as IC for individual cages.

### Genotypes

In this population, 2,374 birds were genotyped using the 600K Affymetrix^®^ Axiom^®^ HD genotyping array [15]. Blood samples were collected from the brachial veins of the animals and DNA was extracted. For the first two generations, all male candidates were genotyped by Ark-Genomics (Edinburgh, UK) during the research project UtopIGe [16]. From the G2 generation onward, male and female reproducers were genotyped at the high-throughput genotyping platform Gentyane (Clermont-Ferrand, France) (cf figure 1).

Each animal was genotyped for 580,961 SNP markers. According to the fifth annotation release of Gallus gallus genome [17], these SNPs were distributed over macro-chromosomes (1 to 5), intermediate chromosomes (6 to 10), micro-chromosomes (11 to 28 and 33), one linkage group (LGE64), two sexual chromosomes Z and W, and a group of 3,724 SNPs with unknown locations.

Genotypes were filtered through four successive steps: individuals with a call rate <95% were removed; SNPs with a MAF <0.05 were removed; SNPs with a call rate <95% were removed; SNPs whose genotype frequencies deviated significantly from the Hardy-Weinberg equilibrium (P < 10^−4^) were removed. Animals showing pedigree incompatibilities were also removed (12 individuals excluded). Thus, 302,102 SNPs (cf table 1) and a total of 1,214 genotyped males and 1,148 genotyped females were kept for the study.

**Table 1.** Summary of the different steps of SNPs quality control

| Genotypes filtration | Number of SNP removed |
| --- | --- |
| MAF ($<0.05$) | 258 772 |
| SNP Call Rate ($<0.95$) | 7 549 |
| Hardy-Weinberg equilibrium ($< 10^{-4}$) | 12 538 |
| SNP retained for analyses | 302 102 |

### Traits

In this paper, traits are named according to Animal Trait Ontology for Livestock [18]. Five egg quality traits related to egg shell quality and internal egg quality, were studied: egg weight (EW), egg shell color (ESC), egg shell strength (ESS), albumen height (AH) and egg shell shape index (ESshape). Summary statistics on traits are given in tables 2 & 3.

**Table 2.**
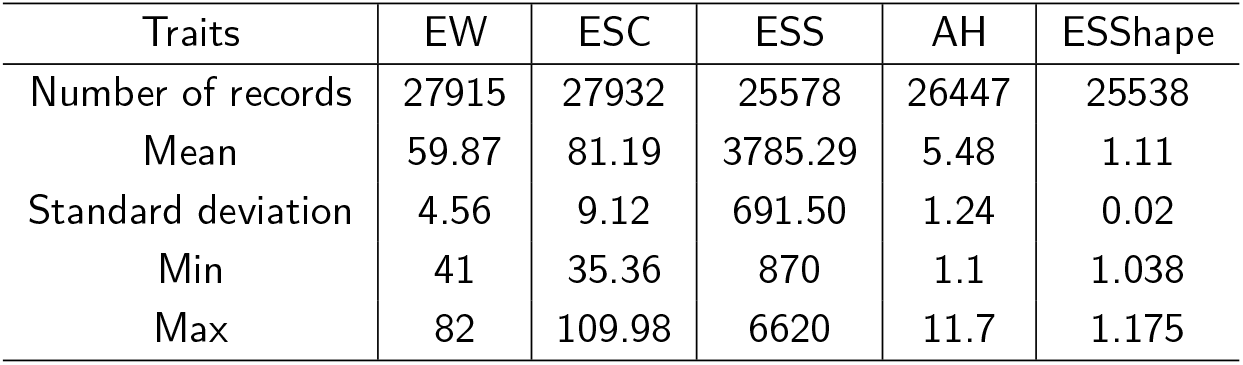
Summary statistics on phenotypic data in CC

#### Trait measurements

At 30 and 50 weeks of age for CC, and once a week for IC, the eggs produced on the farm were collected and quality traits were measured by the company Zootests (Ploufragan, France). The first step consisted in measuring egg short length (SLE in mm) and EW (in g), before calculating ESshape as: 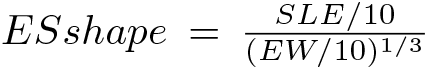. Then, egg shell color was measured using a Minolta chromameter and three traits were recorded: redness of egg shell a^*^, yellowness of egg shell b^*^ and lightness of egg shell L^*^. Egg shell color was then calculated as: *ESC* = 100 *−* (*L*^*^*− a*^*^*− b*^*^). Thirdly, shell strength was measured using a compression machine to evaluate the static stiffness of the shell. Eggs were compressed between two flat plates moving at constant speed. ESS is the maximum force recorded before eggshell fracture (in N, multiplied by 100). Finally, the egg was broken and AH (in mm) was measured using a tripod.

#### Adjustment for environmental effects

For each trait, egg measurements were adjusted for environmental effects using the SAS^®^ 9.4 GLM procedure, based on the following unitrait linear model

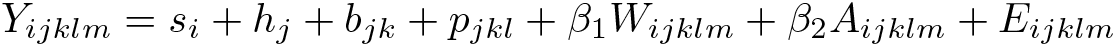

Where *Y_ijklm_* is trait value; *s_i_* is the fixed effect of sire i (514 levels in CC, 421 in IC); *h_j_* is the fixed effect of the hatch j (12 levels); *b_jk_* and *p_jkl_* define the cage location in the poultry house within hatch, respectively the fixed effect of battery *b_jk_* (20 levels in CC, 33 in IC) and the fixed effect of the cage position along the battery *p_jkl_* (58 levels in CC, 65 in IC); *W_ijklm_* is the waiting time between sample and egg measurement covariable (in days) and *A_ijklm_* is the age of the hen covariable (in weeks); *E_ijklm_* is the random residual variable.

For all the traits, the effects of this model were below the significance level (*P <* 0.15), which means that they could be kept into the model. Raw data were then adjusted using the estimates of all effects, except for the sire effect. Extreme values were deleted. Were considered as extreme the values presenting a deviation from the mean higher than 5 phenotypic standard deviations. Finally, a total of more than 25,500 records for CC and more than 65,800 records for IC were retained (cf tables 2 and 3).

**Table 3.**
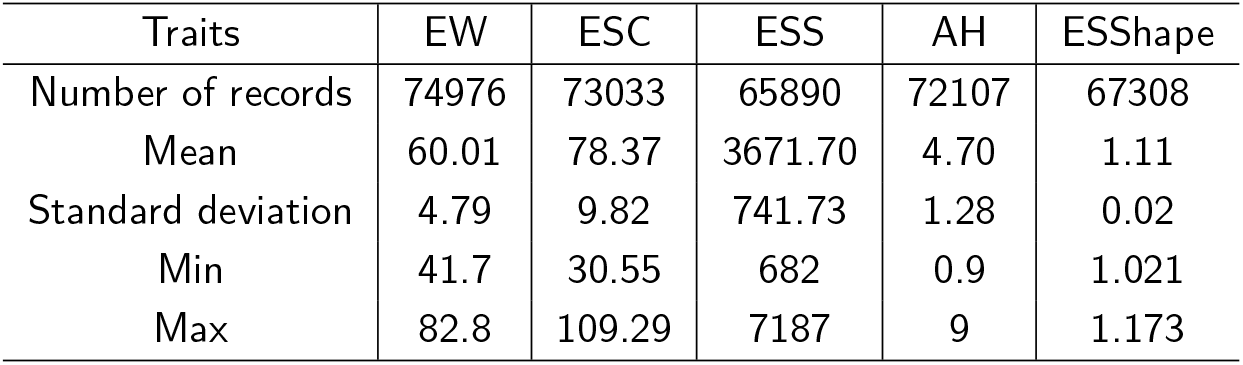
Summary statistics on phenotypic data in IC

### Genetic and genomic evaluations

Performances were centered and standardized before evaluation. Multi-trait evaluations were performed on the five traits, using BLUP (EBV) and single-step GBLUP (GEBV) methodologies [19, 20]. To perform those evaluations, the BLUPF90 family of programs [21] was used. Variance-covariance matrices were estimated using REMLF90. The standard errors of genetic parameters estimates were then obtained with AIREMLF90.

#### Statistical model

The statistical model was the same for all the traits and took into account the fixed and covariables environmental effects described before, plus the random genetic effect of the animal. For CC evaluations, each egg measured was associated with one cage of five full sisters, without knowing which hen laid which egg. As the measurements were repeated twice in CC, each hen had two measured eggs in expectation, but it was not possible to take into account the common environmental effect of the hen. The heritability estimate was calculated as the ratio between the animal variance and the sum of animal and residual variances. Conversely, for IC evaluations, several measurements were available for each hen, and a random common environmental effect of the hen was taken into account in the model. The heritability coefficient was therefore estimated as the ratio between the animal variance and the sum of animal, common environmental and residual variances, and the repeatability coefficient as the ratio between the sum of animal and common environmental variances and the sum of animal, common environmental and residual variances.

#### Candidate populations

To assess the relevance of genetic evaluation versus genomic evaluation, the estimated breeding values, EBV and GEBV, of the selection candidates were compared. In the present study, three different candidate populations were used, two composed of males and one composed of females.

The first male candidate population (C1) was comprised of birds from generation G1 that had daughters (G2) and grand daughters (G3) with performances. This group was made up of 174 sires in CC, and 78 sires in IC. A second male candidate population (C2) was considered in order to increase in the size of the reference population, therefore going from three hatches for C1 to five hatches for C2, which led to a total of 175 sires in CC, and 93 in IC.

In both C1 and C2, the number of sires in IC was smaller than in CC. This is due to the selection carried out before moving them from CC to IC and to the fact that in IC, only sires having at least 8 daughters with performances were used.

In addition to these male populations, a female candidate population (F) was formed using genotyped hens from G2. This group was comprised of 442 females in CC and then moved to IC. The difference between this population and the male ones, was that in IC females had performances available for evaluation and had few daughters (1.6 daughter/hen on average). As opposed to males, females were genotyped starting from G2, which means that is was not possible to have two female candidate populations.

#### Reliability of prediction

To assess the reliability of genetic evaluation and genomic evaluation, the estimated breeding values EBVs and GEBVs had to be compared to the true breeding values (TBVs) of candidates. However, TBV is never known when working with real data set. Moreover, it could not be approximated using Daughter Yield Deviation (DYD) [22], since our candidates had very few offsprings.

Therefore, to estimate the accuracy and bias of prediction of our evaluations, the LR method [23] was used. This method compares evaluation results based on both complete and partial data sets, since the amount of change expected in consecutive genetic evaluations was described as a function of their respective accuracy [24].

All available phenotypes, from G0 to G3, made up the complete data set. Two cases of partial data sets were studied, based on the amount of phenotypic information available when the evaluation was carried out:

- Case 1: The evaluation was carried out at candidates birth, without considering the performances of their contemporary relatives nor the performances of the candidates, in the case of females. The phenotyped population was limited to the ancestors of the candidates.
- Case 2: The evaluation was carried out at 60 weeks of age for CC and at 80 weeks of age for IC, without considering the performances of the progeny of candidates. The phenotyped population included ancestors, contemporary relatives and performances of the candidates, in the case of females. This case corresponds to the scheme classically used in layers selection.

Moreover, for C1 male candidates, the potential gain was also assessed taking into account the performances of their grand-daughters. In that case, the phenotypes of G3 hens were removed so as to obtain a partial data set (case 3).

The LR method relies on three statistics to estimate the accuracy and biases of an evaluation:

- The correlation between (G)EBVs from complete and partial evaluations to estimate relative accuracy
- The difference of means between (G)EBVs from complete and partial evaluations to estimate biases
- The slope of the linear regression of (G)EBVs from complete evaluation on (G)EBVs from partial evaluation, to estimate the over or under dispersion of estimates

To compare genetic evaluation and genomic evaluation, using the same amount of data, a fourth statistics was used: the ratio between the relative accuracy of EBVs and the relative accuracy of GEBVs. This statistics allows to quantify the increase in accuracy expected when moving from genetic evaluation to genomic evaluation.

Furthermore, the significativity of the differences between relative accuracies, e.g. correlations as defined above, was assessed using the Hotelling-Williams test [25]. This test is used to compare two dependent correlations that share a common variable. The null hypothesis means that the two compared correlations are equal. The test statistics under null hypothesis follows the Student law at n-3 degrees of freedom, n being the number of observations. Observed correlations were compared two by two, for EBVs and GEBVs, at a significance threshold of 5%.

## Results

### Genetic parameters

Heritabilities remained steady whether REML was carried out with BLUP or with GBLUP and whichever the data set under analysis, e.g. complete or partial (cases 1,2 and 3). These results were observed whatever the trait or age (CC or IC). Differences ranged from 0% for ESShape in IC to 5% for EW in IC, and values were always higher with GBLUP (data not shown) than with BLUP. Similar results were obtained for repeatabilites. Genetic correlations were even more stable than heritabilities or repeatabilities. Consequently, variance-covariance matrices were set for the rest of the study. The variance-covariance matrix obtained with genomic evaluation throught the use of the complete data set, i.e. REML carried out with the maximum amount of information, was used to perform subsequent BLUP and GBLUP.

Estimates of genetic parameters are given in table 4 for traits measured in CC and in table 5 for traits measured in IC. EW, ESC and ESshape were highly heritable while heritability (resp. repeatability) of ESS and AH were more moderate. For all the traits, heritability was higher in CC than in IC, and was of the same order of magnitude as repeatability in IC. Genetic correlation between EW and AH was positive and moderate, and showed no significant difference between CC and IC. Genetic correlations between ESshape and ESS, or between ESshape and AH, were also positive and moderate, but significantly higher in IC. Genetic correlation between AH and ESS was slighlty lower. EW and ESshape were weakly correlated while ESC was not correlated with other traits in CC but weakly correlated with ESS and ESshape in IC.

**Table 4.** Genetic parameters for CC. In diagonal: trait heritability; upside diagonal: genetic correlations; downside diagonal: phenotypic correlations. In parentheses are standard errors.

| Traits | EW | ESC | ESS | AH | ESShape |
| --- | --- | --- | --- | --- | --- |
| EW | 0.64 (0.02) | 0.02 (0.02) | 0.00 (0.02) | 0.37 (0.02) | 0.12 (0.02) |
| ESC | 0.01 (0.01) | 0.59 (0.02) | -0.02 (0.02) | 0.03 (0.02) | -0.03 (0.02) |
| ESS | -0.02 (0.01) | 0.05 (0.01) | 0.27 (0.02) | 0.12 (0.01) | 0.28 (0.02) |
| AH | 0.17 (0.01) | 0.02 (0.01) | -0.00 (0.01) | 0.34 (0.01) | 0.20 (0.02) |
| ESShape | 0.02 (0.01) | -0.06 (0.01) | 0.18 (0.01) | 0.12 (0.01) | 0.48 (0.02) |

**Table 5.** Genetic parameters for IC. In diagonal: trait heritability/repeatability; upside diagonal: genetic correlations; downside diagonal: phenotypic correlations. In parentheses are standard errors.

| Traits | EW | ESC | ESS | AH | ESShape |
| --- | --- | --- | --- | --- | --- |
| EW | 0.45/0.69 (0.00) | 0.02 (0.01) | 0.14 (0.01) | 0.35 (0.01) | 0.12 (0.01) |
| ESC | 0.02 (0.05) | 0.40/0.57 (0.00) | 0.16 (0.01) | 0.08 (0.01) | -0.16 (0.01) |
| ESS | 0.02 (0.05) | 0.08 (0.03) | 0.23/0.37 (0.03) | 0.21 (0.00) | 0.45 (0.01) |
| AH | 0.10 (0.04) | 0.02 (0.03) | 0.01 (0.00) | 0.26/0.36 (0.01) | 0.41 (0.01) |
| ESShape | -0.03 (0.02) | -0.07 (0.01) | 0.16 (0.01) | 0.10 (0.02) | 0.42/0.56 (0.00) |

### (G)EBVs relative accuracy for male candidates

#### CC traits

As expected, relative accuracy estimates (cf table 6) significantly increased with the amount of phenotypic information available, from case 1 to case 3. In case 1, relative accuracy estimates were not homogeneous and varied depending on the trait. Accuracies for EW and ESshape tended to be low, with a relative accuracy between 0.22 (EW genetic C2) and 0.24 (ESshape genetic C2), even though these traits were more heritable than ESS or AH, for which the relative accuracy was around 0.32 (AH genetic C2). These differences were less present in case 2 and no longer existed in case 3.

**Table 6.**
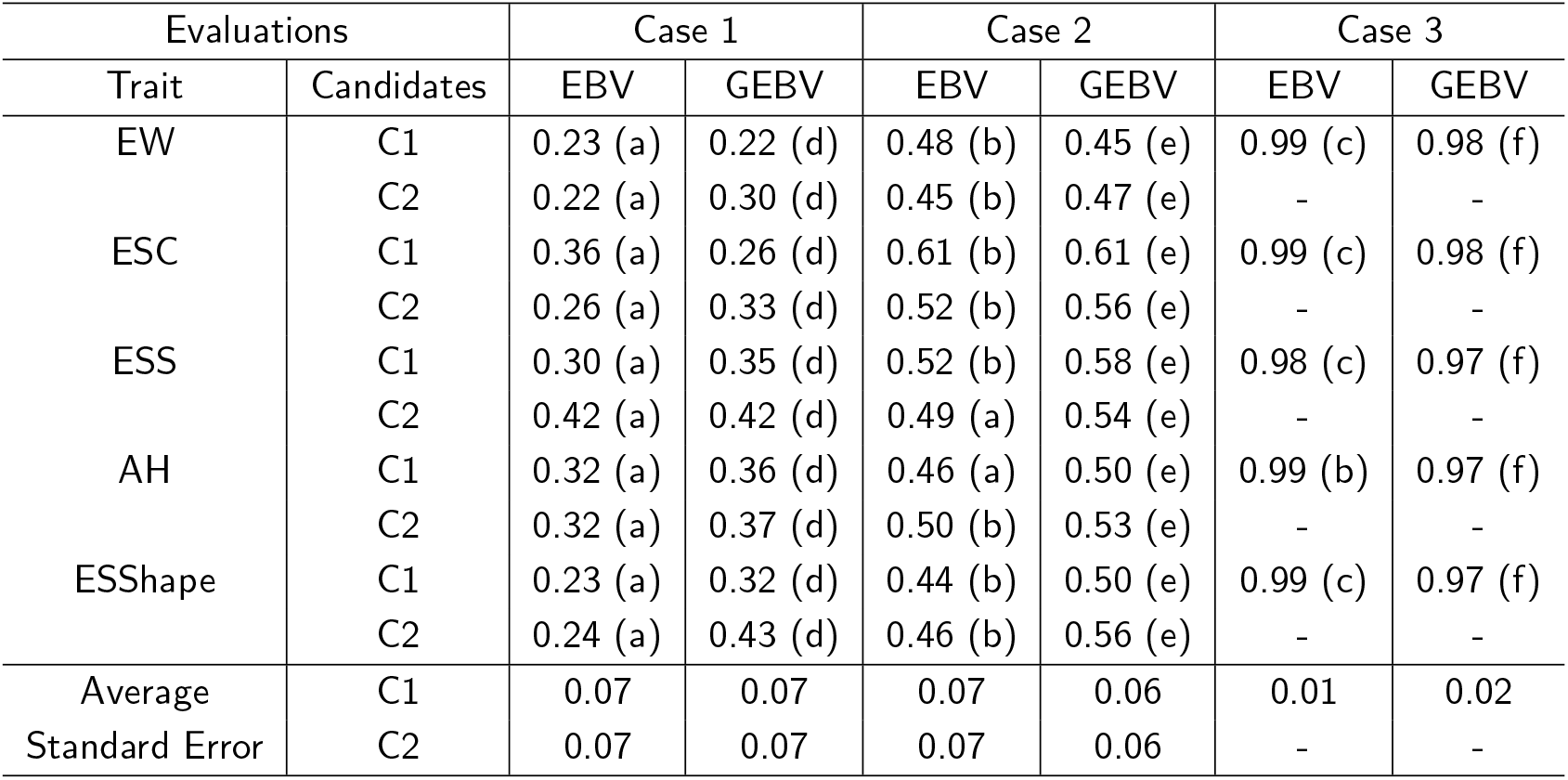
Estimates of accuracy of males (G)EBVs on CC traits. In parentheses are groups determined by an Hotelling-Williams test, at a confidence level of 95%. Groups a, b, and c are used when EBVs correlations are compared to other EBVs correlations; and d, e, f when GEBVs correlations are compared to other GEBVs.

The results of the comparison between genetic evaluation and genomic evaluation are showed in table 7, which gives the ratio of the relative accuracy of BLUP and GBLUP, for each case studied. A value of 1 indicates no difference, values below 1 indicates that genomic evaluation is more accurate, while values above 1 indicates that genetic evaluation is more accurate.

**Table 7.** Ratio of relative accuracy from BLUP to GBLUP, on CC traits for male candidates

| Trait | Candidates | Case 1 | Case 2 | Case 3 |
| --- | --- | --- | --- | --- |
| EW | C1 | 1.05 | 1.05 | 1.05 |
|  | C2 | 0.75 | 0.96 | - |
| ESC | C1 | 1.39 | 1.00 | 1.01 |
|  | C2 | 0.78 | 0.92 | - |
| ESS | C1 | 0.87 | 0.89 | 1.01 |
|  | C2 | 0.99 | 0.90 | - |
| AH | C1 | 0.90 | 0.93 | 1.02 |
|  | C2 | 0.87 | 0.95 | - |
| ESShape | C1 | 0.73 | 0.87 | 1.01 |
|  | C2 | 0.56 | 0.82 | - |

Evaluations carried out at birth (case 1) were the ones showing the greatest amount of difference between genetic evaluation and genomic evaluation. Results proved highly trait-dependent and ranged from 1.39 (ESC C1), e.g. a 39% gain in accuracy with genetic evaluation, compared to genomic evaluation, to 0.56 (ESshape C2), e.g. a 44% gain in accuracy with genomic evaluation, compared to genetic evaluation. There were strong disparities between C1 results, with a mean of 0.99, and C2 results, with a mean of 0.79. Overall, in this case, genomic evaluation allowed for greater accuracy, which can be explained by the size of the reference population. This advantage of genomic evaluation over genetic evaluation was also trait-dependent.

In the evaluations carried out at the age of 18 months (case 2), the differences between genetic evaluation and genomic evaluation were less significant, with values ranging from 1.05 (EW C1) to 0.82 (ESshape C2), and a global mean of 0.93. Differences between C1, with a mean of 0.95, and C2, with a mean of 0.91, were not as significant as they were in case 1, but still existed. Like in case 1, the use of GBLUP allowed for a relative increase in accuracy.

Evaluations carried out in case 3 showed little difference between BLUP and GBLUP, with a mean close to 1.

#### IC traits

As was the case with CC, accuracy estimations were different for each trait (cf table 8), depending on the evaluation scenario. The evolution of accuracy was also linked to the amount of phenotypic information available.

**Table 8.**
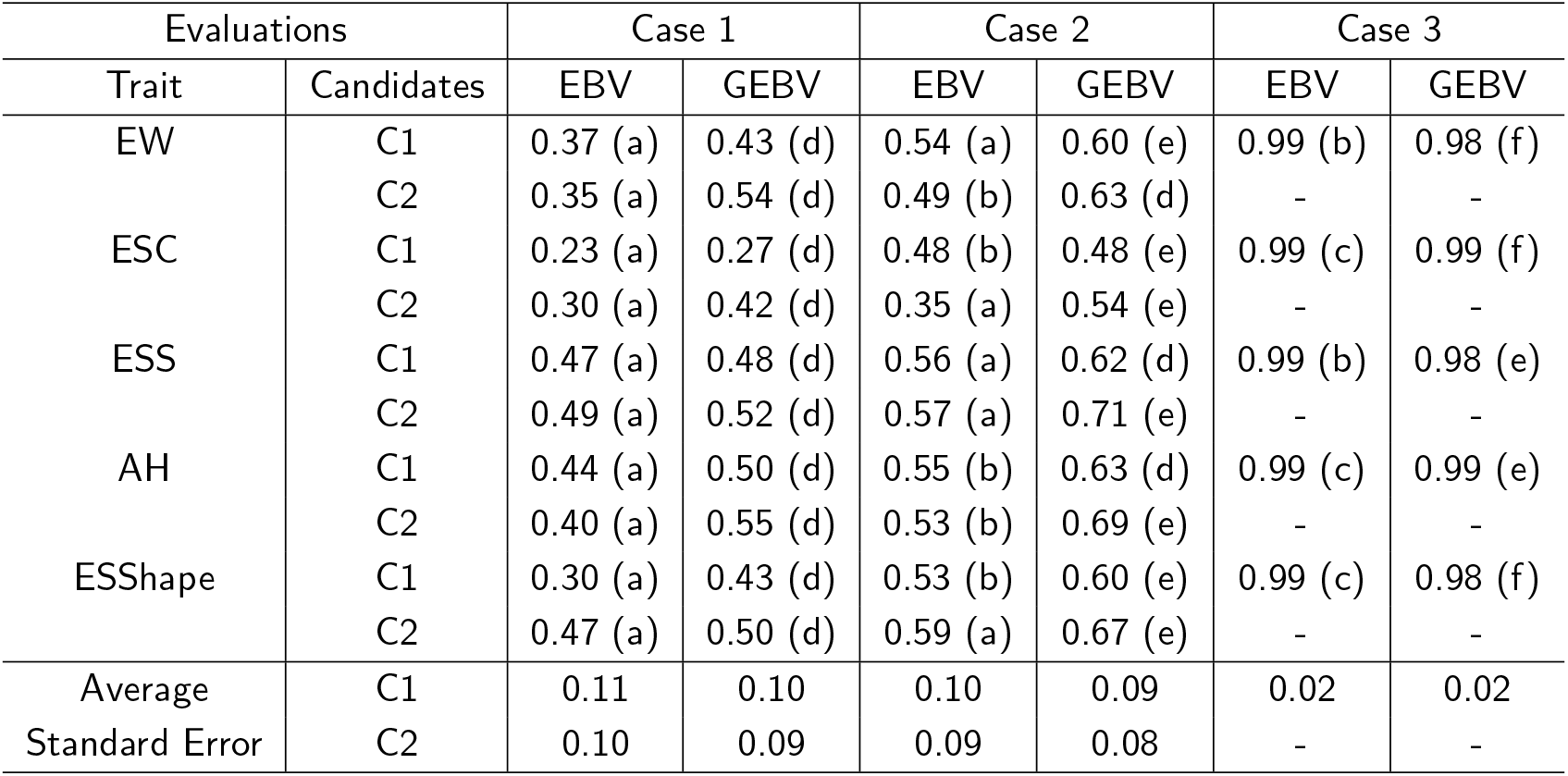
Estimates of accuracy of males (G)EBV on IC traits. In parentheses are groups determined by an Hotelling-Williams test, at a confidence level of 95%. Groups a, b, and c are used when EBVs correlations are compared to other EBVs correlations; and d, e, f when GEBVs correlations are compared to other GEBVs.

Here again, the increase in relative accuracy with genomic evaluation, compared to genetic evaluation, was observed in case 1 and in case 2 (cf table 9). This increase was more significant in IC traits than it was in CC traits, both in case 1 (mean = 0.82) and case 2 (mean = 0.84). The global gain in accuracy observed in evaluations carried out on C2, in comparison to those carried out on C1, was similar to the gain noticed for CC traits.

**Table 9.** Ratio of relative accuracy from BLUP to GBLUP, on IC traits for male candidates

| Trait | Candidates | Case 1 | Case 2 | Case 3 |
| --- | --- | --- | --- | --- |
| EW | C1 | 0.85 | 0.89 | 1.01 |
|  | C2 | 0.65 | 0.78 | - |
| ESC | C1 | 0.84 | 0.99 | 1.01 |
|  | C2 | 0.72 | 0.65 | - |
| ESS | C1 | 0.98 | 0.90 | 1.01 |
|  | C2 | 0.95 | 0.80 | - |
| AH | C1 | 0.87 | 0.88 | 1.01 |
|  | C2 | 0.73 | 0.77 | - |
| ESShape | C1 | 0.70 | 0.88 | 1.00 |
|  | C2 | 0.95 | 0.88 | - |

Evaluations carried out in case 3, showed no differences between BLUP and GBLUP, with a mean very close to 1, as was the case for CC.

### (G)EBVs biases and dispersion for male candidates

The bias statistics has an expected value of 0 if evaluation is unbiased. In both CC and IC (cf tables 10 and 11), biases were low and most often negative, indicating an underestimation of (G)EBVs when using partial data sets. The biases increased as the amount of phenotypic information decreased, from about 0 in case 3 to −0.11 in case 1. Biases were slightly higher with genomic values, compared to genetic values, in any given trait situation. The differences between traits or between candidate population C1 and candidate population C2 varied, without any clear tendency being observed.

**Table 10.** Biases of males (G)EBVs for CC traits

| Evaluations |  | Case 1 |  | Case 2 |  | Case 3 |  |
| --- | --- | --- | --- | --- | --- | --- | --- |
| Trait | Candidates | EBV | GEBV | EBV | GEBV | EBV | GEBV |
| EW | C1 | 0.01 | -0.05 | -0.02 | -0.07 | 0.0 | 0.0 |
|  | C2 | -0.03 | -0.08 | -0.02 | -0.05 | - | - |
| ESC | C1 | -0.04 | -0.11 | 0.01 | -0.07 | 0.01 | -0.01 |
|  | C2 | 0.07 | 0 | 0.01 | -0.01 | - | - |
| ESS | C1 | -0.02 | -0.05 | -0.0 | -0.03 | -0.0 | -0.01 |
|  | C2 | 0.03 | 0 | 0 | -0.02 | - | - |
| AH | C1 | 0.02 | -0.02 | -0.02 | -0.03 | -0.0 | -0.01 |
|  | C2 | 0 | -0.02 | -0.01 | -0.02 | - | - |
| ESShape | C1 | -0.01 | -0.02 | -0.01 | -0.02 | -0.0 | 0.0 |
|  | C2 | 0.01 | 0 | -0.01 | 0 | - | - |

**Table 11.** Biases of males (G)EBVs for IC traits

| Evaluations |  | Case 1 |  | Case 2 |  | Case 3 |  |
| --- | --- | --- | --- | --- | --- | --- | --- |
| Trait | Candidates | EBV | GEBV | EBV | GEBV | EBV | GEBV |
| EW | C1 | 0.02 | 0.08 | 0.0 | 0.05 | -0.0 | 0.0 |
|  | C2 | 0.02 | 0.01 | 0.03 | -0.01 | - | - |
| ESC | C1 | 0.02 | -0.04 | 0.01 | -0.04 | -0.01 | -0.03 |
|  | C2 | 0.07 | -0.01 | 0.03 | -0.02 | - | - |
| ESS | C1 | 0.02 | -0.0 | -0.01 | 0.01 | -0.01 | -0.01 |
|  | C2 | -0.02 | -0.05 | -0.01 | -0.02 | - | - |
| AH | C1 | -0.03 | -0.0 | -0.02 | -0.01 | 0.0 | -0.0 |
|  | C2 | -0.02 | -0.03 | -0.02 | -0.04 | - | - |
| ESShape | C1 | -0.02 | 0.03 | -0.02 | 0.03 | -0.02 | 0.01 |
|  | C2 | -0.01 | -0.01 | -0.01 | -0.01 | - | - |

Unbiased estimators are supposed to have a regression slope equal to 1. This is what was observed in case 3, with regression coefficients estimated between 0.94 and 1.01 in CC and between 0.97 and 1.01 in IC, in the case of genetic evaluation as well as the case of genomic evaluation (tables 12 and 13). In both CC and IC, the slopes decreased below 1 every time the amount of phenotypic information decreased. There was no significant difference between genetic evaluation and genomic evaluation: if slopes were closer to 1 when using genetic evaluation on CC traits, it was quite the opposite in the case of IC traits. Conversely, dispersion appeared significantly higher in CC than in IC: in the case of IC, slopes remained above 0.7, with few exceptions, even in case 1, while they decreased often below 0.7, in the case of CC. The slopes were also strongly linked to the evaluated traits, regardless of the candidate population.

**Table 12.** Slope of regression for males (G)EBVs for CC traits

| Evaluations |  | Case 1 |  | Case 2 |  | Case 3 |  |
| --- | --- | --- | --- | --- | --- | --- | --- |
| Trait | Candidates | EBV | GEBV | EBV | GEBV | EBV | GEBV |
| EW | C1 | 0.53 | 0.41 | 0.69 | 0.61 | 0.98 | 0.97 |
|  | C2 | 0.56 | 0.59 | 0.72 | 0.68 | - | - |
| ESC | C1 | 0.77 | 0.47 | 0.88 | 0.78 | 1.0 | 0.97 |
|  | C2 | 0.59 | 0.50 | 0.69 | 0.67 | - | - |
| ESS | C1 | 0.45 | 0.48 | 0.68 | 0.69 | 0.97 | 0.94 |
|  | C2 | 0.66 | 0.61 | 0.66 | 0.64 | - | - |
| AH | C1 | 0.77 | 0.66 | 0.66 | 0.65 | 0.97 | 0.95 |
|  | C2 | 0.82 | 0.75 | 0.86 | 0.83 | - | - |
| ESShape | C1 | 0.43 | 0.53 | 0.71 | 0.69 | 0.99 | 0.98 |
|  | C2 | 0.55 | 0.76 | 0.79 | 0.81 | - | - |

**Table 13.** Slope of regression of (G)EBVs for males evaluation on IC traits

| Evaluations |  | Case 1 |  | Case 2 |  | Case 3 |  |
| --- | --- | --- | --- | --- | --- | --- | --- |
| Trait | Candidates | EBV | GEBV | EBV | GEBV | EBV | GEBV |
| EW | C1 | 0.72 | 0.73 | 0.76 | 0.78 | 0.99 | 0.97 |
|  | C2 | 0.72 | 0.89 | 0.9 | 0.89 | - | - |
| ESC | C1 | 0.45 | 0.50 | 0.82 | 0.80 | 0.99 | 1.02 |
|  | C2 | 0.54 | 0.57 | 0.54 | 0.66 | - | - |
| ESS | C1 | 1.07 | 0.90 | 0.82 | 0.81 | 1.04 | 1.02 |
|  | C2 | 0.91 | 0.75 | 0.84 | 0.83 | - | - |
| AH | C1 | 0.90 | 0.96 | 0.97 | 1.00 | 1.00 | 0.97 |
|  | C2 | 0.82 | 0.94 | 0.97 | 0.98 | - | - |
| ESShape | C1 | 0.55 | 0.54 | 0.93 | 0.82 | 1.01 | 0.99 |
|  | C2 | 0.89 | 0.61 | 1.10 | 0.98 | - | - |

### (G)EBVs relative accuracy for females

#### CC traits

As was the case for males, accuracy estimations of (G)EBVs for females were not homogeneous, depending on the trait (table 14): some traits were evaluated with greater accuracy than others, and accuracy evolution was not the same for all the traits, depending on the scenario. However, these differences were not the same than those noticed with males. Relative accuracy was nonetheless generally higher for females, especially in case 2 where females had their performances taken into account.

**Table 14.**
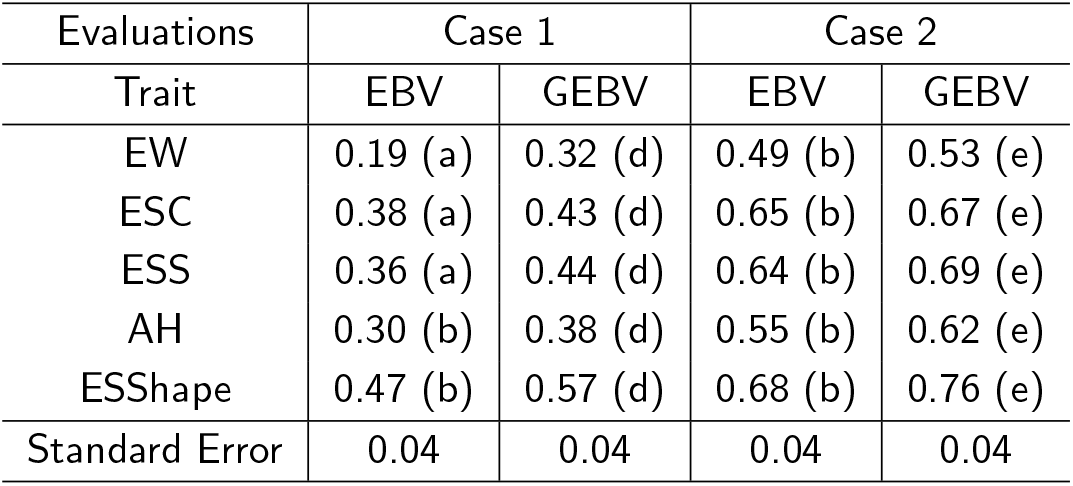
Estimates of accuracy of females (G)EBVs on CC traits. In parentheses are groups determined by an Hotelling-Williams test, at a confidence level of 95%. Groups a, and b are used when EBVs correlations are compared to other EBVs correlations; and d,e when GEBVs correlations are compared to other GEBVs.

Furthermore, genomic values were always more accurate than genetic values and, like with males, the gain increased when the amount of phenotypic information was low (table 15). Indeed, evaluations carried out at birth (case 1), showed a significant increase in accuracy with GBLUP evaluation, compared to BLUP evaluation. The mean of ratios was close to 0.78, i.e. a 22% gain in accuracy with genomic evaluation, compared to genetic evaluation. Regarding evaluations carried oud in case 2, where the performances of the females were taken into account, this value was only 0.92, an 8% gain in accuracy.

**Table 15.**
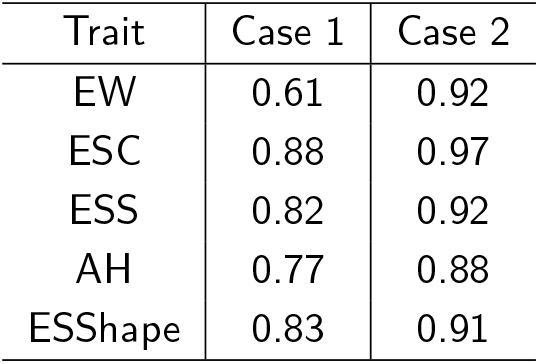
Ratio of the relative accuracy from BLUP to GBLUP, on CC traits for female candidates

| Trait | Case 1 | Case 2 |
| --- | --- | --- |
| EW | 0.61 | 0.92 |
| ESC | 0.88 | 0.97 |
| ESS | 0.82 | 0.92 |
| AH | 0.77 | 0.88 |
| ESShape | 0.83 | 0.91 |

#### IC traits

As was the case for CC, correlations were moderate (cf table 16) and varied depending on the trait in case 1, while they were always very high (with a minimum of 0.93) in case 2, where the performances of the females were taken into consideration.

**Table 16.** Estimates of accuracy of females (G)EBVs on IC traits. In parentheses are groups determined by an Hotelling-Williams test, at a confidence level of 95%. Groups a, and b are used when EBVs correlations are compared to other EBVs correlations; and c,d when GEBVs correlations are compared to other GEBVs.

| Evaluations | Case 1 |  | Case 2 |  |
| --- | --- | --- | --- | --- |
| Trait | EBV | GEBV | EBV | GEBV |
| EW | 0.34 (a) | 0.44 (d) | 0.94 (b) | 0.93 (e) |
| ESC | 0.46 (a) | 0.54 (d) | 0.97 (b) | 0.96 (e) |
| ESS | 0.30 (a) | 0.44 (d) | 0.94 (b) | 0.94 (e) |
| AH | 0.35 (a) | 0.45 (d) | 0.96 (b) | 0.96 (e) |
| ESShape | 0.50 (a) | 0.60 (d) | 0.98 (b) | 0.97 (e) |
| Standard Error | 0.04 | 0.04 | 0.01 | 0.01 |

In case 1, the increase in accuracy noticed with GBLUP evaluation, in comparison to BLUP evaluation (table 17) was of the same order of magnitude than it was for CC, with a mean of ratios close to 0.79. In case 2, this value was between 1.00 and 1.01, depending on the trait.

**Table 17.** Ratio of the relative accuracy from BLUP to GBLUP, on IC traits for female candidates

| Trait | Case 1 | Case 2 |
| --- | --- | --- |
| EW | 0.77 | 1.01 |
| ESC | 0.85 | 1.01 |
| ESS | 0.69 | 1.01 |
| AH | 0.79 | 1.00 |
| ESShape | 0.83 | 1.01 |

### (G)EBVs biases and dispersion for female candidates

As was the case for males, and both in CC and IC (cf tables 18 and 19), biases were low and often negative (with an exception in case 1 EW and ESC). This showed an under estimation of (G)EBVs when evaluating with partial data sets. A similar increase in biases was observed in females when the amount of phenotypic information decreased. Here again, no clear relationship could be noticed between traits and biases, nor between the type of evaluation carried out (genetic or genomic) and biases, contrary to what was observed with males.

**Table 18.** Biases of females (G)EBVs for CC traits

| Evaluations | Case 1 |  | Case 2 |  |
| --- | --- | --- | --- | --- |
| Trait | EBV | GEBV | EBV | GEBV |
| EW | 0.13 | 0.09 | 0 | 0.01 |
| ESC | 0.10 | 0.02 | 0.01 | -0.04 |
| ESS | 0 | -0.03 | 0 | -0.01 |
| AH | 0.01 | -0.01 | 0 | -0.01 |
| ESShape | -0.04 | -0.04 | 0 | -0.01 |

**Table 19.** Biases of females (G)EBVs for IC traits

| Evaluations | Case 1 |  | Case 2 |  |
| --- | --- | --- | --- | --- |
| Trait | EBV | GEBV | EBV | GEBV |
| EW | 0.04 | 0.06 | 0 | -0.01 |
| ESC | 0.08 | 0 | -0.01 | -0.04 |
| ESS | 0.02 | 0.01 | -0.01 | -0.01 |
| AH | 0.02 | 0.02 | 0 | 0 |
| ESshape | 0.03 | 0.03 | -0.02 | -0.01 |

As to what regards regression coefficient, both in CC and IC (cf tables 20 and 21), the results were similar to those observed using male candidates. Except for ESshape in case 1, the regression coefficients decreased below 1 every time the amount of phenotypic information decreased. As was the case for males, there was not any significant difference between genetic evaluation and genomic evaluation, and dispersion seemed higher for CC than for IC. Once again, slopes were strongly linked to the trait evaluated, in any given case.

**Table 20.**
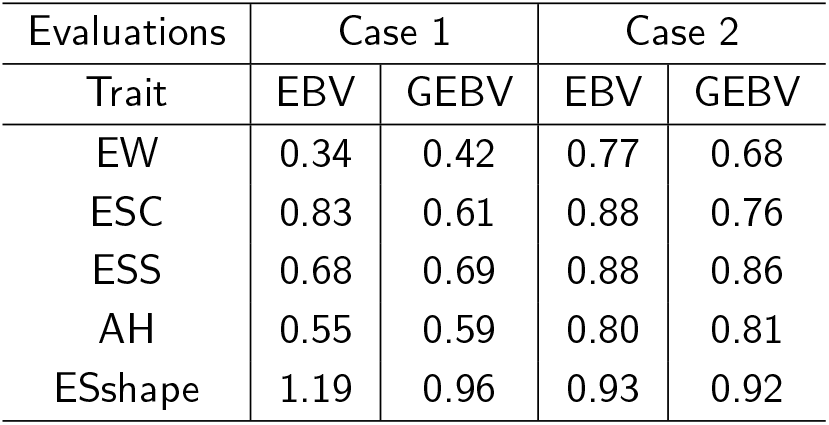
Slope of regression for females (G)EBVs for CC traits

| Evaluations | Case 1 |  | Case 2 |  |
| --- | --- | --- | --- | --- |
| Trait | EBV | GEBV | EBV | GEBV |
| EW | 0.34 | 0.42 | 0.77 | 0.68 |
| ESC | 0.83 | 0.61 | 0.88 | 0.76 |
| ESS | 0.68 | 0.69 | 0.88 | 0.86 |
| AH | 0.55 | 0.59 | 0.80 | 0.81 |
| ESshape | 1.19 | 0.96 | 0.93 | 0.92 |

**Table 21.**
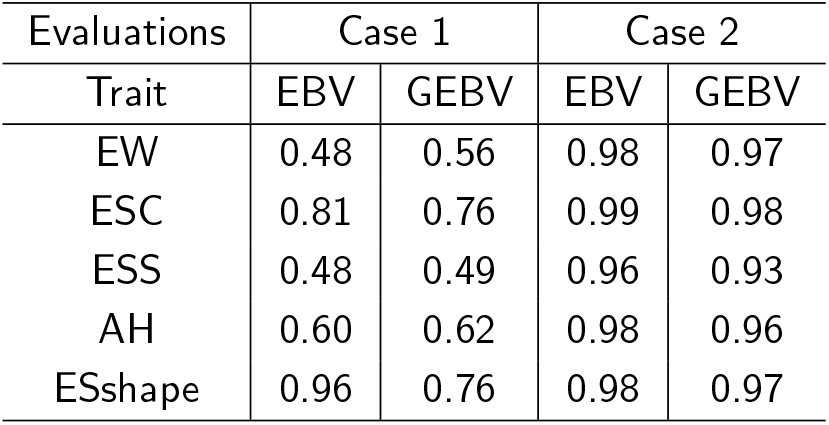
Slope of regression for females (G)EBVs for IC traits

| Evaluations | Case 1 |  | Case 2 |  |
| --- | --- | --- | --- | --- |
| Trait | EBV | GEBV | EBV | GEBV |
| EW | 0.48 | 0.56 | 0.98 | 0.97 |
| ESC | 0.81 | 0.76 | 0.99 | 0.98 |
| ESS | 0.48 | 0.49 | 0.96 | 0.93 |
| AH | 0.60 | 0.62 | 0.98 | 0.96 |
| ESshape | 0.96 | 0.76 | 0.98 | 0.97 |

## Discussion

### Genetic parameters

Estimates of heritability, repeatability and genetic correlations (cf tables 4 for CC and 5 for IC) were in accordance with the literature [26, 4]. Moderate to high heritability coefficients allow to expect significant genetic gains through selection on the five egg quality traits. Selection carried out in order to increase egg shell strength should lead to an increase in albumen height, which is favorable to a combined selection of these traits. It should also cause an increase in egg weight, which can be potentially favorable or unfavorable, depending on the breeding goal of the line, as well as an increase in egg short length at a given weight, which is unfavorable, e.g. the egg would be less ovoid. Finally, egg shell color is lightly, but favorably, correlated to other traits.

### Relevance of genomic evaluation for male candidates

In case 1 and in case 2, the results generally highlighted a greater accuracy of the evaluation of male candidates with GBLUP than with BLUP, in any given scenario, and particularly for what regards IC traits. The difference between CC results and IC results can be explained by the nature of data, as indeed IC data, referring to the hen itself and not just to the full sisters, allowed for the construction a more accurate evaluation model. Otherwise, the results obtained in case 3 showed that the information about the grand daughters had little impact on the evaluation. The fact of not using the performances of the grand-daughters does not seem to have any direct negative impact on evaluation accuracy.

The results observed in scenarios using C2 candidates tend to confirm those obtained using C1 candidates, and sometimes even amplified them. The difference between C1 results and C2 results could be explained by the increase in the size of the reference population, which went from 3 batches for C1 to 5 batches for C2, still with the same number of candidates. This increase in the size of the reference population had an impact on evaluation, with more candidate haplotypes represented into the reference population, as shown by Rabier et *al* [9].

### Relevance of genomic evaluation for female candidates

The differences between traits, and the relationship between phenotypic information and accuracy, noticed with males were also observed with females. Genomic evaluation provided more accurate evaluations than genetic evaluation, excepted for case 2, where IC traits were used. GEBV accuracy was capped at around 0.70 in CC, while it came close to 1 in IC. This difference was due to the fact that in CC, phenotypes are not related to a single bird, but to a cage of full sisters, which means that instead of being performed on the female, the evaluation was performed on its family.

Evaluations performed on a candidate population of females in IC (case 2) showed a relative accuracy of (G)EBV, which came close to 1. This result was due to the fact that the performances of the females were taken into account for the evaluation: the addition of genomic information did not increase the gain in accuracy. Furthermore, as opposed to male candidates, females had very few daughters with performances: the lack of information about the performances of the daughters had little impact on the estimate of the value of the females.

## Conclusion

The results of the present study lead to several conclusions regarding the use of genomic evaluation of egg quality traits in a layers line.

In both CC and IC, the reliability of genetic evaluation or of genomic evaluation varied greatly depending on the trait. This heterogeneousness in the impact on the evaluation can be explained by the differences found in the genetic architecture of the traits [16].

As to what regards the comparison between genomic evaluation and genetic evaluation, it was noticed that genomic evaluation most of the time proved more accurate than genetic evaluation, with the same amount of phenotypic information used. It was also observed that the increase in accuracy of genomic evaluation was higher when phenotypic information was restricted, as genotypes partly compensated for the loss of phenotypic information.

Regarding the size of the reference population, it was observed that adding a generation, from C1 to C2, had an effect on the evaluation, as Weng and al. showed [13]. An augmentation of the reference population, from 1 to 2 generations, increased evaluation accuracy, especially when there was little phenotypic information available.

These results can be interesting for the poultry industry. The reduction of evaluation accuracy could be compensated for by an increase in selection pressure and a shorter generation interval, especially for males. This would allow for an optimization of the expected genetic gain.

Finally, for what regards egg quality traits and as far as males are concerned, it seems winning to move from a selection at 18 months of age to a selection at birth. Indeed, for some traits, there would be a significant loss in evaluation accuracy in the case of CC, although the loss would remain acceptable for IC. Depending on the weight of each trait in the breeding goal, this strategy would allow for a significant genetic gain on the global objective, through an increment in selection pressure and a reduction in the generation interval, on the male pathway. It would therefore be feasible to increase the number of male candidates for selection from 200 to 2,000, and to reduce the generation interval from 18 months to 6 months.

The results also highlighted the fact that, for females, switching from a selection at 18 months of age to a selection at birth would result in a significant loss in evaluation accuracy. In the case of females, selection pressure should not be increased that much. However, the generation interval could be reduced from 18 to 6 months here as well. Unlike in the case of males, this strategy needs to be studied in more details for females, in order to assess whether the implementation of genomic evaluation at birth would be an interesting option.

## Ethics statement

All information used in this study were collected from hens raised in the context of layers selection. These animals and the scientific investigations described herein are therefore not to be considered as experimental animals per se, as defined in EU directive 2010/63 and subsequent national application texts. Consequently, we did not seek ethical review and approval of this study as regarding the use of experimental animals. All animals were reared on the Novogen nucleus herd in compliance with national regulations pertaining to livestock production and according to procedures approved by the French Veterinary Services.

## Availability of data and materials

The datasets used and/or analysed throughout the present study are available from the corresponding author on reasonable request.

## Author’s contributions

DPD performed the statistical analyses and drafted the manuscript. FlH filtered the genotype data. AVa and TB supervised animal management and production. PLR supervised the analyses and participated in drafting the manuscript. PLR and TB conceived the study. All authors contributed to the ideas and methods. All authors read and approved the final manuscript.

## Acknowledgements

This research project was partly supported by the French national research agency ?Ç£ANR?Ç within the framework of project ANR-10-GENOM BTV-015 UtOpIGe. DPD is a PhD fellow supported by the Brittany region (France) and the INRA metaprogram SelGen.

## References

1 Meuwissen TH, Hayes BJ, Goddard ME. Prediction of total genetic value using genome-wide dense marker maps. Genetics. 2001 Apr;157(4):1819–1829. Available from: http://www.ncbi.nlm.nih.gov/pmc/articles/PMC1461589/.

2 Hayes BJ, Bowman PJ, Chamberlain AJ, Goddard ME. Invited review: Genomic selection in dairy cattle: progress and challenges. J Dairy Sci. 2009 Feb;92(2):433–443.

3 Wolc A, Kranis A, Arango J, Settar P, Fulton JE, O’Sullivan NP, et al. Implementation of genomic selection in the poultry industry. ResearchGate. 2016 Jan;6(1):23. Available from: https://www.researchgate.net/publication/289494029_Implementation_of_genomic_selection_in_the_poultry_industry.

4 Wolc A, Zhao HH, Arango J, Settar P, Fulton JE, O’Sullivan NP, et al. Response and inbreeding from a genomic selection experiment in layer chickens. Genetics Selection Evolution. 2015;47:59. Available from: http://dx.doi.org/10.1186/s12711-015-0133-5.

5 Liu T, Qu H, Luo C, Shu D, Wang J, Lund MS, et al. Accuracy of genomic prediction for growth and carcass traits in Chinese triple-yellow chickens. BMC Genetics. 2014 Oct;15:110. Available from:https://doi.org/10.1186/s12863-014-0110-y.

6 Daetwyler HD, Villanueva B, Woolliams JA. Accuracy of predicting the genetic risk of disease using a genome-wide approach. PLoS ONE. 2008;3(10):e3395.

7 Liu Z, Seefried FR, Reinhardt F, Rensing S, Thaller G, Reents R. Impacts of both reference population size and inclusion of a residual polygenic effect on the accuracy of genomic prediction. Genet Sel Evol. 2011 May;43(1):19. Available from: http://www.ncbi.nlm.nih.gov/pmc/articles/PMC3107172/.

8 Erbe M, Hayes BJ, Matukumalli LK, Goswami S, Bowman PJ, Reich CM, et al. Improving accuracy of genomic predictions within and between dairy cattle breeds with imputed high-density single nucleotide polymorphism panels. J Dairy Sci. 2012 Jul;95(7):4114–4129.

9 Rabier CE, Barre P, Asp T, Charmet G, Mangin B. On the Accuracy of Genomic Selection. PLOS ONE. 2016 Jun;11(6):e0156086. Available from: http://journals.plos.org/plosone/article?id=10.1371/journal.pone.0156086.

10 Elsen JM. Approximated prediction of genomic selection accuracy when reference and candidate populations are related. Genet Sel Evol. 2016 Mar;48. Available from: http://www.ncbi.nlm.nih.gov/pmc/articles/PMC4778372/.

11 Clark SA, Hickey JM, Daetwyler HD, van der Werf JH. The importance of information on relatives for the prediction of genomic breeding values and the implications for the makeup of reference data sets in livestock breeding schemes. Genet Sel Evol. 2012 Feb;44(1):4. Available from: http://www.ncbi.nlm.nih.gov/pmc/articles/PMC3299588/.

12 Habier D, Fernando RL, Dekkers JCM. The Impact of Genetic Relationship Information on Genome-Assisted Breeding Values. Genetics. 2007 Dec;177(4):2389–2397. Available from: http://www.genetics.org/content/177/4/2389.

13 Weng Z, Wolc A, Shen X, Fernando RL, Dekkers JCM, Arango J, et al. Effects of number of training generations on genomic prediction for various traits in a layer chicken population. Genetics Selection Evolution. 2016;48:22. Available from: http://dx.doi.org/10.1186/s12711-016-0198-9.

14 Lourenco DaL, Misztal I, Tsuruta S, Aguilar I, Lawlor TJ, Forni S, et al. Are evaluations on young genotyped animals benefiting from the past generations? J Dairy Sci. 2014;97(6):3930–3942.

15 Kranis A, Gheyas AA, Boschiero C, Turner F, Yu L, Smith S, et al. Development of a high density 600K SNP genotyping array for chicken. BMC Genomics. 2013 Jan;14:59.

16 Romé H, Varenne A, Hérault F, Chapuis H, Alleno C, Dehais P, et al. GWAS analyses reveal QTL in egg layers that differ in response to diet differences. Genetics Selection Evolution. 2015 Oct;47:83. Available from: https://doi.org/10.1186/s12711-015-0160-2.

17 Warren WC, Hillier LW, Tomlinson C, Minx P, Kremitzki M, Graves T, et al. A New Chicken Genome Assembly Provides Insight into Avian Genome Structure. G3 (Bethesda). 2017;7(1):109–117.

18 Domaines d’ATOL - Animal Trait Ontology for Livestock;. Available from: http://www.atol-ontology.com/.

19 Aguilar I, Misztal I, Johnson DL, Legarra A, Tsuruta S, Lawlor TJ. Hot topic: a unified approach to utilize phenotypic, full pedigree, and genomic information for genetic evaluation of Holstein final score. J Dairy Sci. 2010 Feb;93(2):743–752.

20 Misztal I, Legarra A, Aguilar I. Computing procedures for genetic evaluation including phenotypic, full pedigree, and genomic information. J Dairy Sci. 2009 Sep;92(9):4648–4655.

21 Misztal I, Tsuruta S, Strabel T, Auvray B, Druet T, Lee D. BLUPF90 and related programs. In: Proceedings of the 7th World Congress on Genetics Applied to Livestock Production. vol. Vol. 28; 2002. p. 743.

22 VanRaden PM. Efficient methods to compute genomic predictions. J Dairy Sci. 2008 Nov;91(11):4414–4423.

23 Legarra A, Reverter A. Semi-parametric estimates of population accuracy and bias of predictions of breeding values and future phenotypes using the LR method. Genet Sel Evol. 2018 Nov;50(1):53.

24 Reverter A, Golden BL, Bourdon RM, Brinks JS. Technical note: detection of bias in genetic predictions2. Journal of Animal Science. 1994 Jan;72(1):34–37. Available from: https://academic.oup.com/jas/article/72/1/34-37/4632556.

25 Van Sickle J. nalyzing correlations between stream and watershed attributes. Journal of the American Water Resources Association 39(3):717–726. 2003;.

26 Beaumont C, Calenge F, Chapuis H, Fablet J, Minvielle F, Tixier-Boichard M. Génétique de la qualité de l’œuf - Inra Prod.Anim., 23(2), 123–132; 2010. Available from: http://www6.inra.fr/productions-animales/2010-Volume-23/Numero-2-2010/Genetique-de-la-qualite-de-l-aeuf.

